# Sperm collection and computer-assisted sperm analysis in the teleost model Japanese medaka (*Oryzias latipes*)

**DOI:** 10.1101/2022.05.24.492481

**Authors:** Lauren Closs, Amin Sayyari, Romain Fontaine

**Author notes:** **Corresponding Author:** Amin Sayyari, Romain Fontaine. **Email Addresses of Co-Authors:** Lauren Closs, Amin Sayyari, Romain Fontaine.

## Abstract

Japanese medaka (*Oryzias latipes*) are a teleost fish and an emerging vertebrate model for ecotoxicology, developmental, genetics, and physiology research. Medaka are also used extensively to investigate vertebrate reproduction, which is an essential biological function as it allows a species to perpetuate. Sperm quality is an important indicator of male fertility and thus reproduction success. Techniques for extracting sperm and sperm analysis are well documented for many species, including for teleost fish. Collecting semen is relatively simple in larger fish but can be more complicated in small model fish as they produce less sperm and are more delicate. This article therefore describes two methods of sperm collection in the small model fish Japanese medaka: testes dissection and abdominal massage. We demonstrate that both approaches are viable for medaka and show that abdominal massage can be performed a repeated number of times as the fish quickly recover from the procedure. We also describe a protocol for computer-assisted sperm analysis in medaka to objectively assess several important indicators of medaka sperm quality (motility, progressivity, duration of motility, relative concentration). The use of these procedures combined with the other advantages of using this small teleost model will greatly improve the understanding of the environmental, physiological, and genetic factors influencing fertility in vertebrate males.

**SUMMARY:** This article describes two quick and efficient methods to collect semen from the small model fish medaka (*Oryzias latipes*), as well as a protocol to reliably assess sperm quality using computer-assisted sperm analysis (CASA).

## INTRODUCTION

Japanese medaka are small egg-laying freshwater teleost fish, native to East Asia. Medaka have become an excellent vertebrate model system for ecotoxicology, developmental genetics, genomics and evolutionary biology and physiology studies (Shima & Mitani, 2004; Wittbrodt et al., 2001). Like the popular zebrafish, they are relatively easy to breed and highly resistant to many common fish diseases (Shima & Mitani, 2004; Wittbrodt et al., 2001). There are several advantages of using medaka as a model, including a short generation time, transparent embryos (Shima & Mitani, 2004; Wittbrodt et al., 2001), and a sequenced genome (Kasahara et al., 2007). Unlike zebrafish, medaka have a sex-determining gene (Matsuda et al., 2002), and a high temperature (from 4–40 °C) and salinity (euryhaline species) tolerance (Sakamoto et al., 2001). Also, many genetic and anatomical tools, as well as protocols (Ager-Wick et al., 2018; Fontaine et al., 2018; Fontaine & Weltzien, 2019; Murata et al., 2019; Porazinski et al., 2010; Royan et al., 2020, 2021), have been developed in medaka to facilitate the study of its biology.

Reproduction is an essential physiological function as it allows a species to perpetuate. Vertebrate reproduction requires a myriad of precisely orchestrated events, including the production of oocytes in females and the production of sperm in males. Sperm are unique cells, produced through the complex process of spermatogenesis, in which there are a number of checkpoints in place to guarantee delivery of a high-quality product (Bhat et al., 2021). Gamete quality has become a focus in aquaculture and fish population studies, due to its impact on fertilization success and larval survival. Sperm quality is therefore an important indicator of male fertility in vertebrates.

Three useful factors for assessing fish sperm quality are motility, progressivity, and longevity. Percent motility and progressive motility are common indicators of sperm quality as progressive motion is necessary for and correlates strongly with fertilization success (Kime et al., 2001; van der Horst et al., 2021). Duration of movement is also an important indicator in fish as sperm remain fully motile for less than 2 minutes in most teleost species and the trajectory of sperm is generally less linear than in mammals (Kime et al., 2001). However, many studies assessing sperm motility in the past relied on subjective or semi-quantitative methods of analyzing sperm (reviewed Kime et al., 2001; Rurangwa et al., 2004). For instance, sperm motility in medaka has been estimated in the past visually under a microscope (Yang & Tiersch, 2009). It has also been estimated by recording sperm movement and using imaging software to merge frames and measure swimming path and velocity (Hara Y et al., 2007; Hashimoto S et al., 2009; Kawana et al., 2003). Such approaches often lack robustness, providing different results according to the person performing the analysis (Gallego et al., 2018; Kime et al., 2001).

Computer-assisted sperm analysis (CASA) was initially developed for mammals. CASA is a fast quantitative method to assess sperm quality by recording and measuring velocity and trajectory in an automated manner (Kime et al., 2001). In fishes, it has been used in different species to monitor the effects of several water pollutants on sperm quality, for identifying interesting progenitors to improve broodstock, to improve the efficiency of cryopreservation and storage, and to optimize conditions for fertilization (Kime et al., 2001). It is therefore a powerful tool for reliably assessing sperm quality in different vertebrate species. However, due to the important diversity in reproductive strategies between fishes, the sperm of teleost fish differs from that of mammals and from one fish species to another. Teleost fish, which primarily fertilize eggs externally by releasing gametes into water, have highly concentrated sperm that are relatively simple in structure with no acrosome, unlike mammals, which fertilize internally and therefore do not have to compensate for dilution in water, but do have to withstand more viscous fluids (van der Horst et al., 2021). Additionally, sperm from most fish move rapidly but are fully motile for less than 2 minutes after activation, although there are several exceptions (Browne et al., 2015; Kime et al., 2001). Because motility can decrease rapidly, extreme care should be taken with the timing of analysis after activation when determining a sperm analysis protocol for fish.

Reproduction is one of the fields in biology where teleosts and medaka have been extensively used as model organisms. Indeed, medaka males show interesting reproductive and social behaviors, such as mate guarding (Arias Padilla et al., 2021; Okuyama et al., 2017). In addition, several transgenic lines exist to study the neuroendocrine control of reproduction in this species (Hodne et al., 2019; Karigo et al., 2014; Okubo et al., 2006). Semen collection, a procedure that is relatively simple in larger fish, can be more complicated in small model fish as they produce less semen and are more delicate. For this reason, many studies involving sperm collection in medaka extract semen by crushing dissected testes (Kowalska A et al., 2007; Kowalska et al., 2011; Yang & Tiersch, 2009). Some studies also use abdominal massage to express the semen directly into activating medium (Hara Y et al., 2007; Hashimoto S et al., 2009; Kawana et al., 2003), however with this method it is difficult to visualize the amount and color of semen extracted. In zebrafish, abdominal massage is commonly used to express semen, which is immediately collected in a capillary tube (Draper BW & Moens CB, 2009; Harvey et al., 1982; Wasden et al., 2017). This method enables estimation of the volume of semen, as well as observation of ejaculate color, which is a quick and simple indicator of sperm quality (Draper BW & Moens CB, 2009; Wasden et al., 2017). Therefore, a clear and well described protocol for sperm collection and analysis is lacking for medaka.

This article therefore describes two different methods of sperm collection in the small model fish Japanese medaka, testes dissection and abdominal massage with capillary tubes. We demonstrate that both approaches are viable for medaka and show that abdominal massage can be performed a repeated number of times as the fish quickly recovers from the procedure. We also describe a protocol for computer-assisted sperm analysis in medaka to reliably measure several important indicators of medaka sperm quality (motility, progressivity, longevity, and relative sperm concentration). The use of these procedures combined with the other advantages of using this small teleost model will greatly improve the understanding of the environmental, physiological, and genetic factors influencing fertility in vertebrate males.

## PROTOCOL

All experimentation and animal handling were conducted in accordance with the recommendations on the experimental animal welfare at Norwegian University of Life Sciences. The experiments were performed using adult male Japanese medaka.

1. **Instruments and solutions preparation**
  1.1. Prepare anesthetic stock solution (0.6% Tricaine).
    1.1.1. Dilute 0.6 g of Tricaine (MS-222) in 100 mL of 10X Phosphate Buffer Saline (PBS).
    1.1.2. Distribute 2 mL of the Tricaine stock solution into several 2 mL plastic tubes and store at -20 °C until use.
  1.2. Prepare recovery water (0.9% sodium chloride [NaCl] solution).
    1.2.1. Add 27 g of NaCl into 3 L of aquarium water.
    1.2.2. Store the solution at room temperature until use.
  1.3. Adjust activation medium if necessary (Hank’s balanced salt solution [HBSS]). NOTE: HBSS can be purchased commercially or made in the laboratory.
    1.3.1. Measure the pH of the HBSS using a pH meter. Adjust the pH if necessary, using hydrochloric acid or sodium hydroxide, so the final pH is 7.1 - 7.3.
    1.3.2. Measure the osmolality of the HBSS using an osmometer for future reporting.
    1.3.3. Store the solution at room temperature until use.
  1.4. Prepare holding sponge
    1.4.1. Cut a soft sponge to fit snugly in a Petri dish.
    1.4.2. Cut a straight line in the middle of the sponge that is long enough to receive the fish (3-4 cm) and about 1 cm deep (**Figure 1A**). This slit in the sponge will hold the fish ventral side up to expose the cloaca.
2. **Sperm collection** NOTE: Sperm collection can be achieved by two different methods: abdominal massage or testes dissection.
  2.1. Sperm collection by abdominal massage
    2.1.1 Prepare 0.03% anesthetic solution by diluting one tube of Tricaine stock (0.6%) in 38 mL of aquarium water in a 100 mL glass container.
    2.1.2 Prepare instruments including blunt end smooth forceps and a 10 µL disposable calibrated glass micropipette and aspirator tube assembly (**Figure 1A**). Dampen the holding sponge prepared in step 1.4, with anesthetic solution.
    2.1.3 Prepare tubes with 36 µL of activating solution for immediate analysis. Preheat the activating solution in a water bath or incubator set to 27 °C, at least 5 minutes. NOTE: When pooling samples from multiple fish, use 36 µL of activating solution per fish.
    2.1.4 Anesthetize the fish by putting it into the anesthetic solution for 30-90 seconds. NOTE: The duration of the anesthesia depends on the size and weight of the fish and must be adapted. To ensure that the fish is fully anesthetized, the fish body can be pinched gently using forceps. If the fish does not react, the massage can be started.
    2.1.5 Take the fish out of the anesthetic solution and use a paper towel or gentle wipe to dry the bottom of the fish. Place the fish in the trough of the damp holding sponge ventral side up, so its gills are exposed to the anesthetic solution in the sponge (**Figure 2A**).
    2.1.6 If the area around the cloaca is wet, gently dry the underside of the fish with a disposable tissue wipe.
    2.1.7 Place the fish in the holding sponge under a dissecting microscope and place the micropipette with aspirator tube attached against the cloaca of the fish (**Figure 2B**).
    2.1.8 Massage the abdomen of the fish by gently squeezing with blunt end smooth forceps in a rostral to caudal motion while simultaneously sucking to collect the expelled semen into the pipette (**Figure 2C**).
    2.1.9 Release the fish from the sponge into recovery water. Allow them to recover in the solution for at least 15 minutes before returning them to the aquarium system.
    2.1.10 Transfer the semen into a tube with preheated activating solution and pipette up and down several times by sucking and blowing on the aspirator tube assembly.
    2.1.11 Homogenize the diluted sperm gently by flicking the tubes before analysis.
  2.2. Sperm collection by testes dissection
    2.2.1 Prepare 0.08% of euthanasia solution by diluting two tubes of Tricaine stock (0.6%) in 26 mL of aquarium water in a 100 mL glass container.
    2.2.2 Prepare dissection tools including blunt and fine forceps, and small dissecting scissors (**Figure 1B**).
    2.2.3 Prepare a tube for each sample with 120 µL of activating solution for immediate analysis. Preheat the activating solution in a water bath or incubator set to 27 °C, at least 5 minutes. NOTE: For pooling samples from multiple fish, use 120 µL of activating solution per fish.
    2.2.4 Euthanize the fish by putting it into the 0.08% anesthetic solution for 30-90 seconds. NOTE: The duration depends on the size and weight of the fish. To ensure that the fish is euthanized, wait for operculum movements to cease. The fish should not react to the touch of forceps.
    2.2.5 Remove the fish from the euthanasia solution and gently dry the fish with a paper towel or gentle wipe. NOTE: At this step, the fish can be weighed to later calculate the gonadosomatic index (GSI, gonadal weight/ body weight).
    2.2.6 Place the fish under a dissecting microscope with its left lateral side facing up (**Figure 3A**).
    2.2.7 Using small dissecting scissors, cut a flap dorsally from the cloaca and then across the ribs to the gills to expose the internal organs (**Figure 3B**).
    2.2.8 Locate the testes, cut the attachment at both ends with fine forceps, and remove the testes (**Figure 3C**). NOTE: To calculate the GSI, the testes can be weighed at this step. Work quickly to avoid drying of the tissue.
    2.2.9 Transfer the testes to a tube with preheated activating solution.
    2.2.10 Use forceps to crush the testes several times against the side of the tube to release the sperm. Sperm release can usually be visualized and will make the solution slightly cloudy.
    2.2.11 Homogenize the diluted sperm gently by flicking the tubes before analysis.
3. **Sperm Analysis with CASA system**
  3.1. The CASA system (SCA Evolution) should be set up according to the manual with a microscope using a green filter and 10X objective with phase contrast.
  3.2. Place a disposable 20 micron counting chamber slide onto a warming plate or in an incubator set to 27 °C, at least 5 minutes.
  3.3. Pipette the sample into the chamber on the slide until it fills the chamber without overfilling. Carefully wipe away excess sample from the entrance of the chamber with a cotton tip or gentle wipe to prevent floating cells. Place the slide under the microscope on a heated stage set to 27 °C and analyze using CASA system.
  3.4. Open the SCA Evolution software and select the motility module.
  3.5. Set the configuration for medaka as shown in **figure 4B**.
  3.6. Select “analyze” to look at the sample under the microscope. NOTE: If the microscope icon is red, the microscope lighting needs to be adjusted for the program to accurately track sperm. Adjust the brightness of the microscope so the tail movement of the sperm is clearly seen. The icon should be blue.
  3.7. Ensure the microscope is focused and select “analyze” to record the sperm in the field. Move the slide so a new area of the sample is in the frame and repeat to capture 3-5 different fields of view. Avoid fields with air bubbles, cell masses, or artifacts.
  3.8. Select “results” to view the results. NOTE: If the fields are outlined in red, follow the system’s prompts to delete the fields that vary too much in concentration or motility.
  3.9. Double click on a field to view the results for the individual field or to manually check for any mislabeled or untracked spermatozoa. Right click on individual spermatozoa to relabel the motility if necessary (**Figure 4A**).

**Figure 1:**
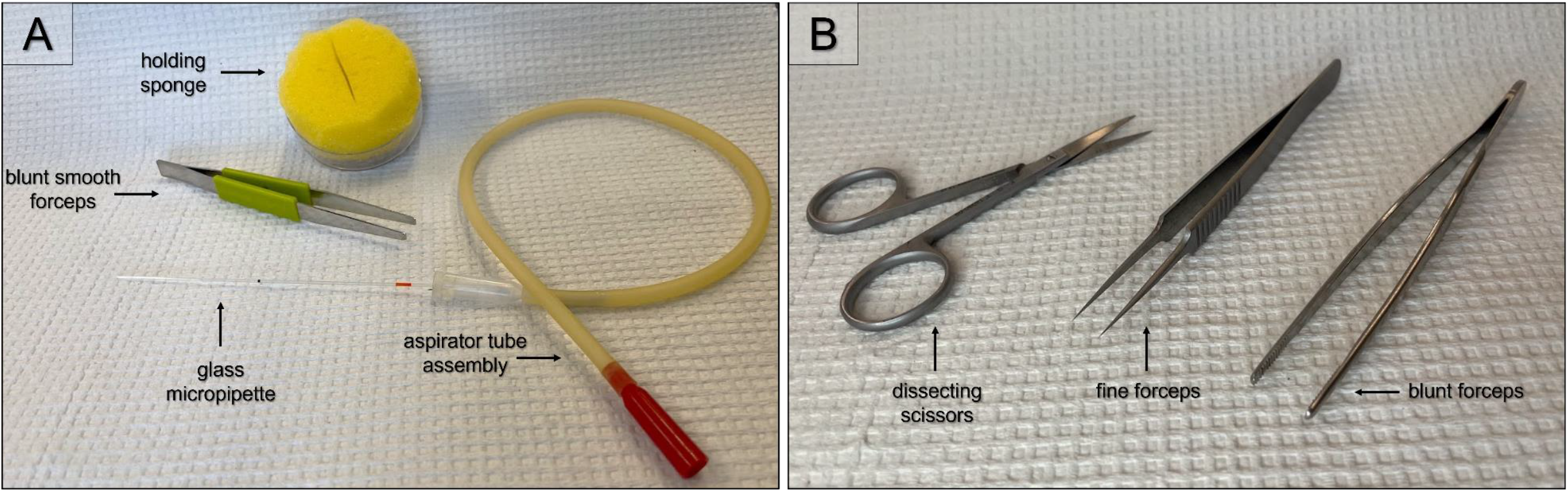
Instruments. **A)** Holding sponge, blunt smooth forceps, and 10 µL disposable calibrated glass micropipette with aspirator tube assembly for semen sampling by abdominal massage; **B)** blunt forceps, fine forceps, and small dissecting scissors for testes sampling.

**Figure 2:**
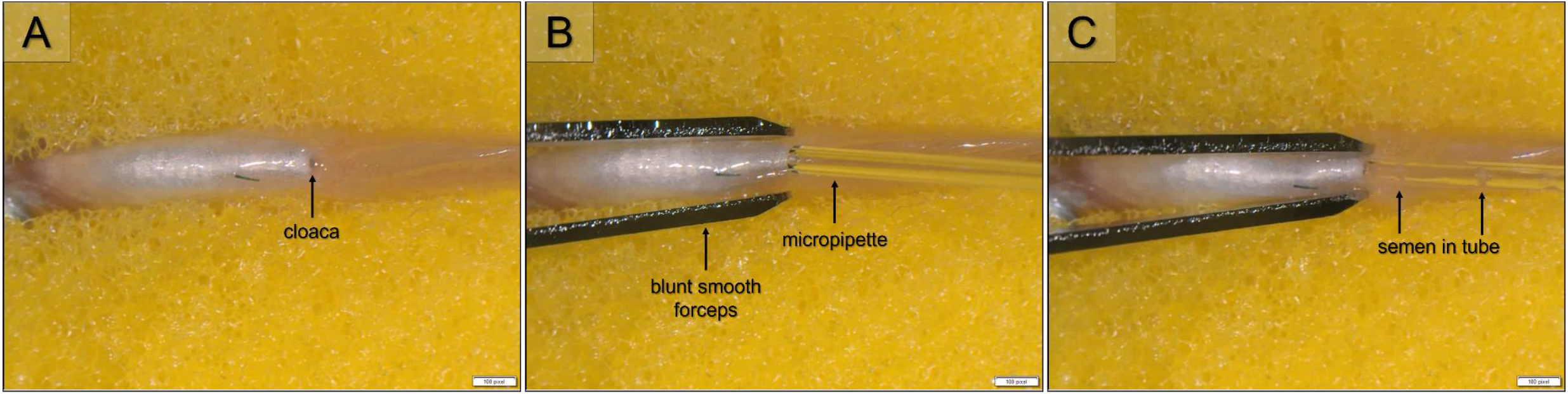
Semen sampling by abdominal massage. **A)** Position of fish in holding sponge, with gills exposed to anesthetic solution in sponge and cloaca facing up; **B)** Position of blunt smooth forceps on abdomen and micropipette against cloaca; **C)** Semen in micropipette after gentle massage and sucking.

**Figure 3:**
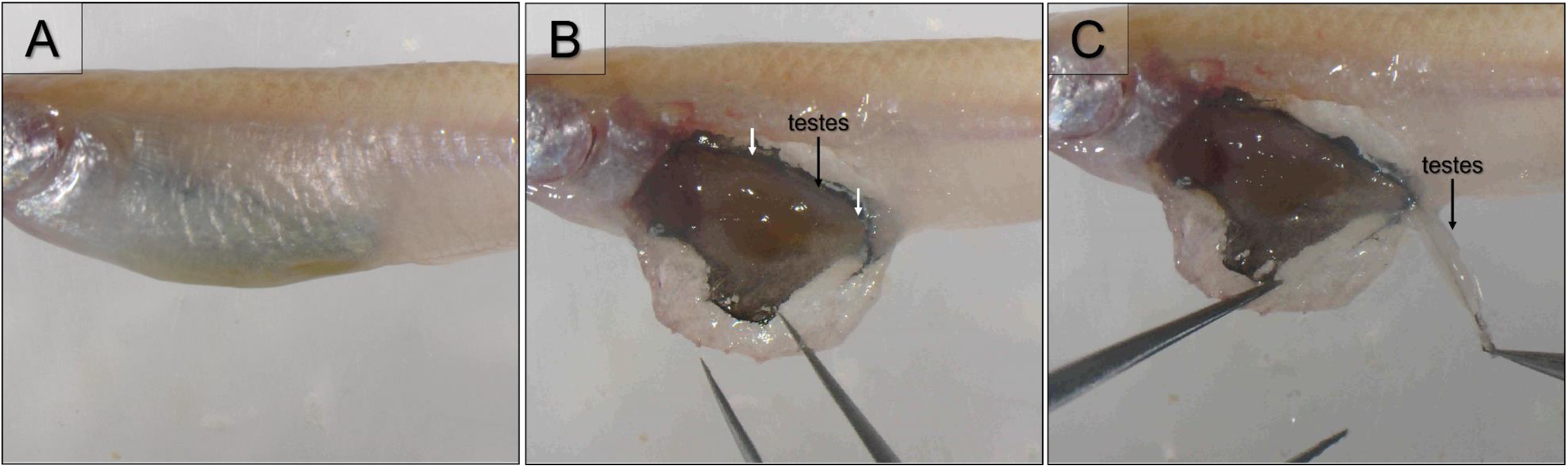
Semen sampling by testes dissection. **A)** Position of fish for testes dissection; **B)** Lateral view of internal organs with white arrows indicating where the testes are attached; **C)** Remove the testes by cutting the attachment at both ends with fine forceps.

**Figure 4:**
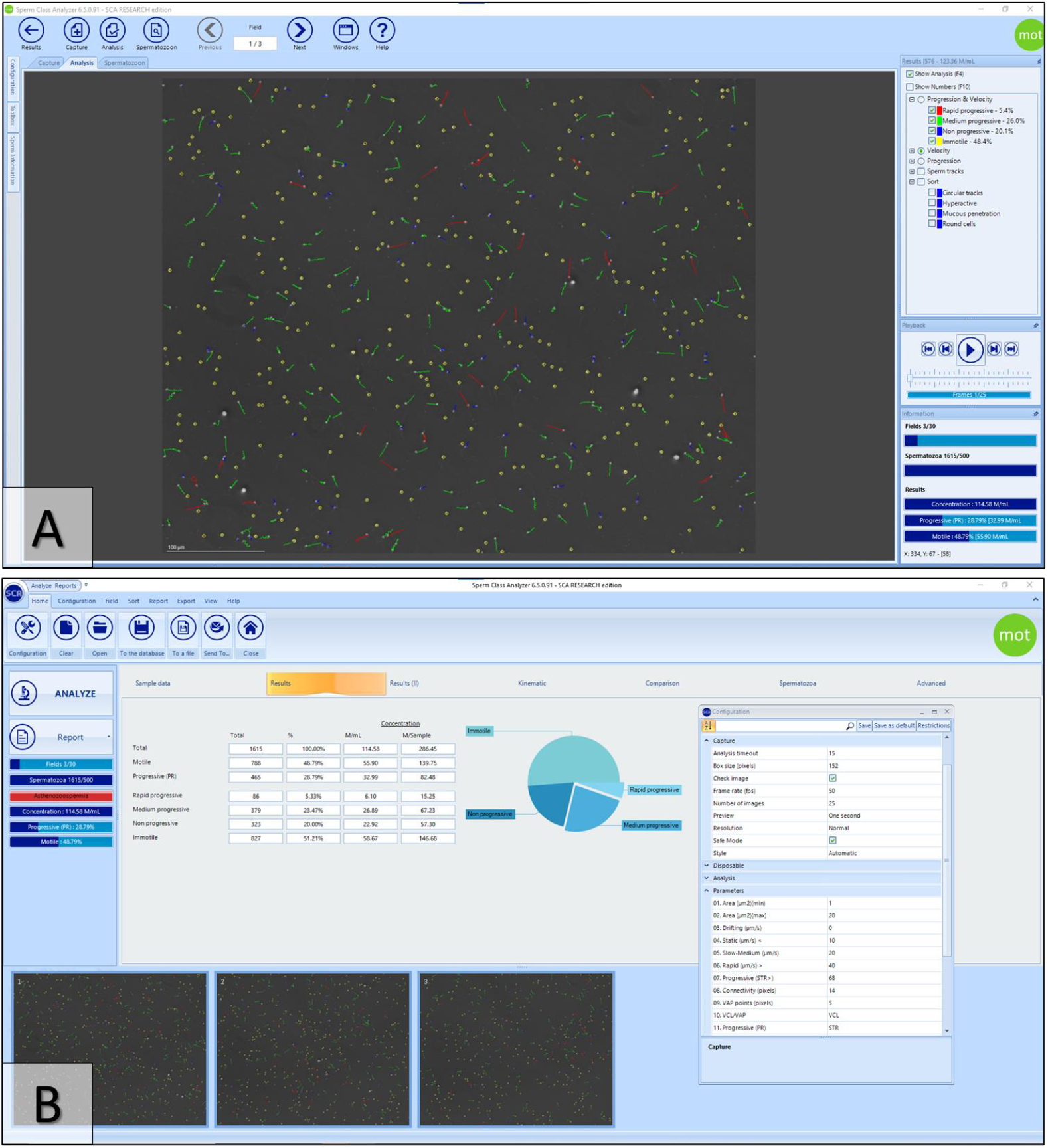
SCA Evolution software screenshot. **A)** Sperm tracking results for one field. View field data on the right side and double click spermatozoa to view individual data; **B)** Results summary for all fields with configuration menu open.

## REPRESENTATIVE RESULTS

### Type of data obtained

Sperm motility analysis from the SCA Evolution software provides data on motility (percentage of motile and immotile sperm), as well as progressivity (percentage of progressive and non-progressive sperm), and velocity (percentage of rapid, medium, and slow-moving sperm). It also combines progressivity and velocity (rapid progressive, medium progressive, non-progressive). These labels are based on measurements (**Figure 5A**) and calculations (**Figure 5B**) of spermatozoon movement, which are provided by the program (**Supplemental Table 1**). For medaka, we adapted the following thresholds from the recommended zebrafish parameters based on previous literature (Acosta et al., 2016; Castellini et al., 2011; Hara Y et al., 2007) and the distribution of our data. Motility is based on curvilinear velocity (VCL) with immotile < 10 µm/s ≤ slow < 20 µm/s ≤ medium ≤ 40 µm/s < rapid. Sperm are considered progressive if straightness index (STR) > 68%.

**Figure 5:**
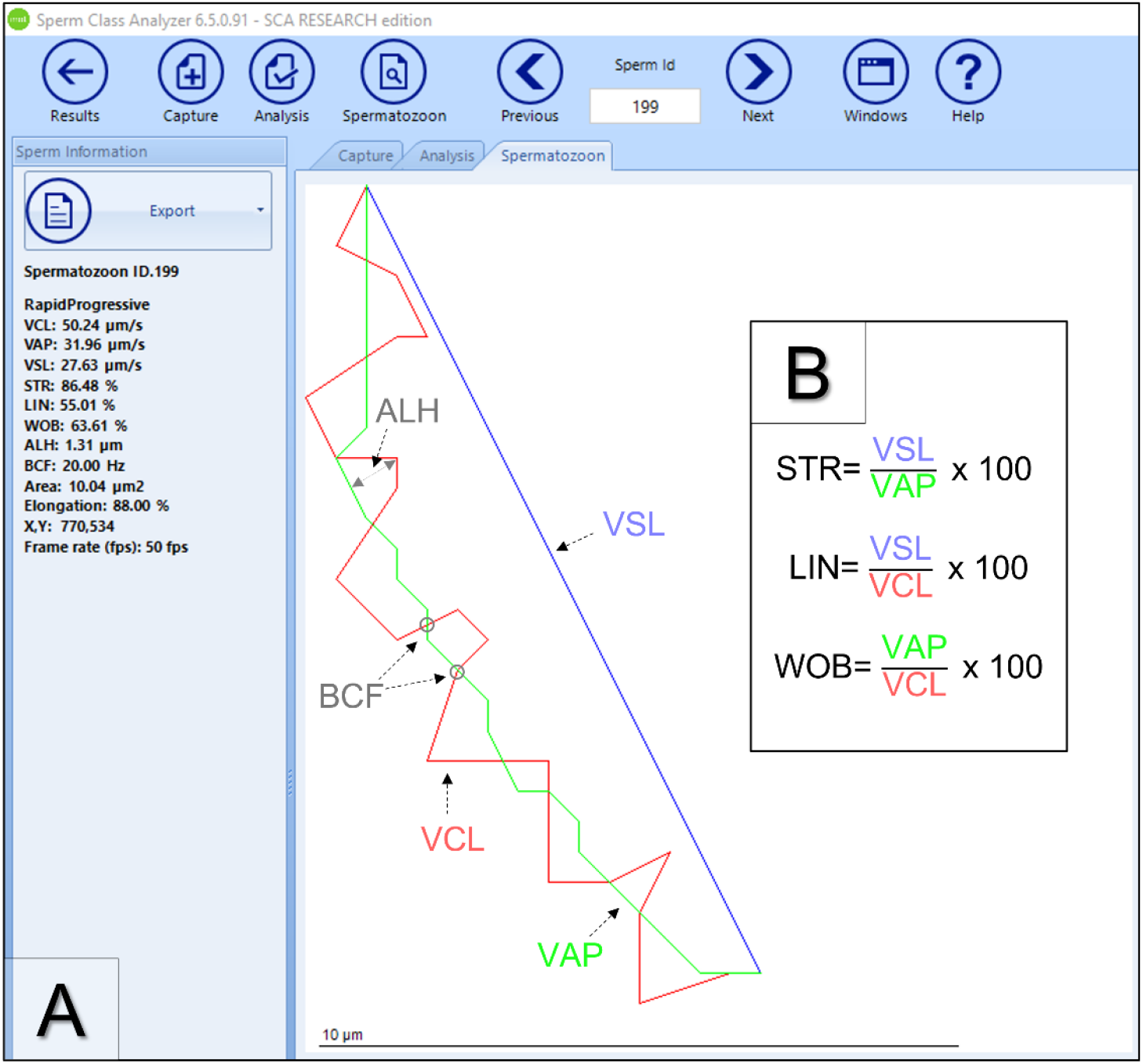
Analysis of spermatozoon movement. **A)** Data recorded by the CASA system include curvilinear velocity (VCL, velocity calculated using the distance along the actual path), average path velocity (VAP, velocity calculated using the distance of the average path), straight line velocity (VSL, velocity calculated using the distance between the start and end of the sperm track), amplitude of lateral head displacement (ALH, magnitude of lateral displacement of a sperm head about its average path), beat cross frequency (BCF, rate at which the curvilinear path crosses the average path); **B)** Calculated values from the CASA system include straightness index (STR, linearity of the average path), linearity index (LIN, linearity of the curvilinear path, and wobble (WOB, oscillation of the actual path about the average path).

SCA Evolution also calculates the concentration of sperm, if provided the volume of the sample and the dilution. While the abdominal massage method is helpful for visualizing the coloration and confirming the presence of the semen, the volume is too small to accurately measure. However, relative concentration to compare fish or treatment groups can be calculated for samples taken by testes dissection, as long as the whole testis is dissected and diluted in the same volume of liquid.

### Sperm motility evaluation

#### Different activating solutions

Surprisingly, sperm sampled by both abdominal massage and testes dissection were immotile in aquarium water (16 mOsmol/kg) (**Figure 6**) and in aquarium water adjusted with NaCl solution to 34 mOsmol/kg. There was also no motility in deionized water (−1 mOsmol/kg), or deionized water adjusted to 23 mOsmol/kg. Sperm were motile in HBSS (287 mOsmol/kg), as well as in HBSS diluted with deionized water to 36 and 113 mOsmol/kg, although the percent motility was significantly reduced at 36 mOsmol/kg (**Figure 7**).

**Figure 6:**
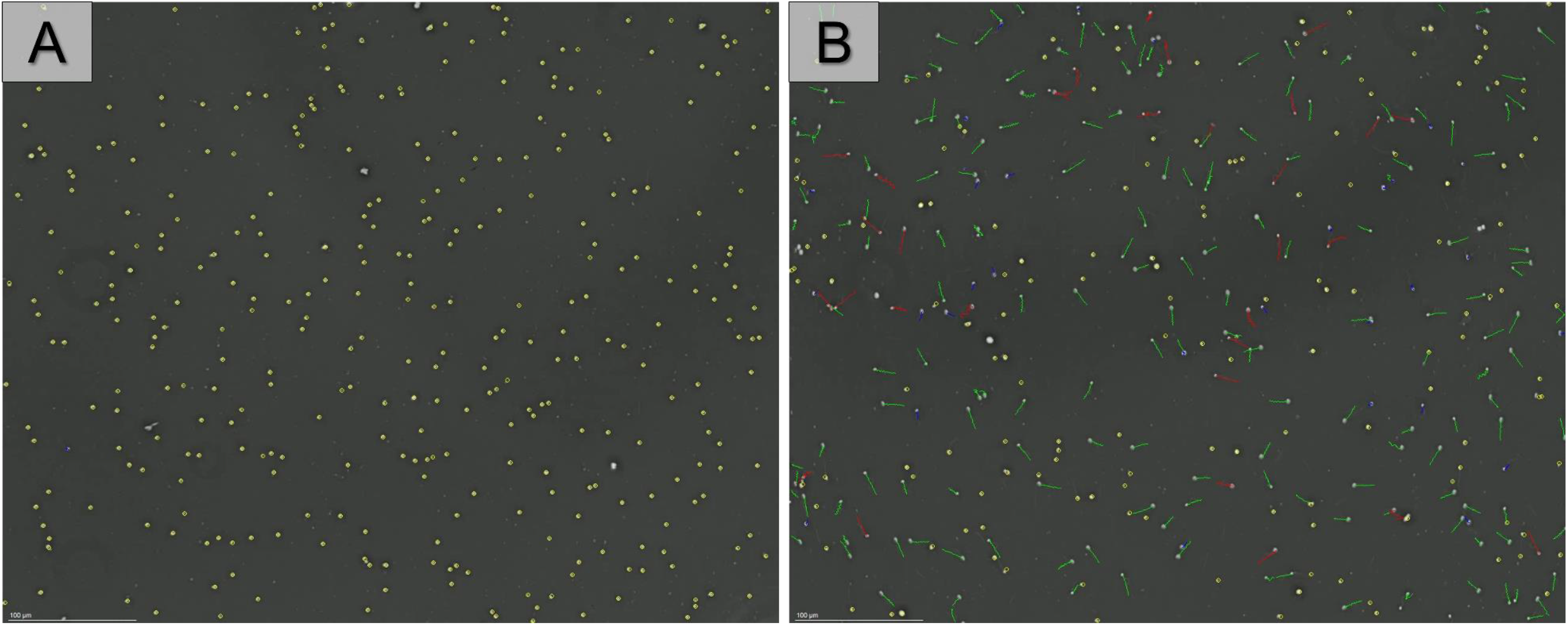
Unsuccessfully and successfully activated sperm tracks. Sperm sampled by testes dissection in **A)** aquarium water (16 mOsmol/kg) and **B)** unadjusted HBSS (287 mOsmol/kg). Yellow circles label immotile sperm and colored lines indicate sperm movement: red (rapid progressive), green (medium progressive), blue (non-progressive).

**Figure 7:**
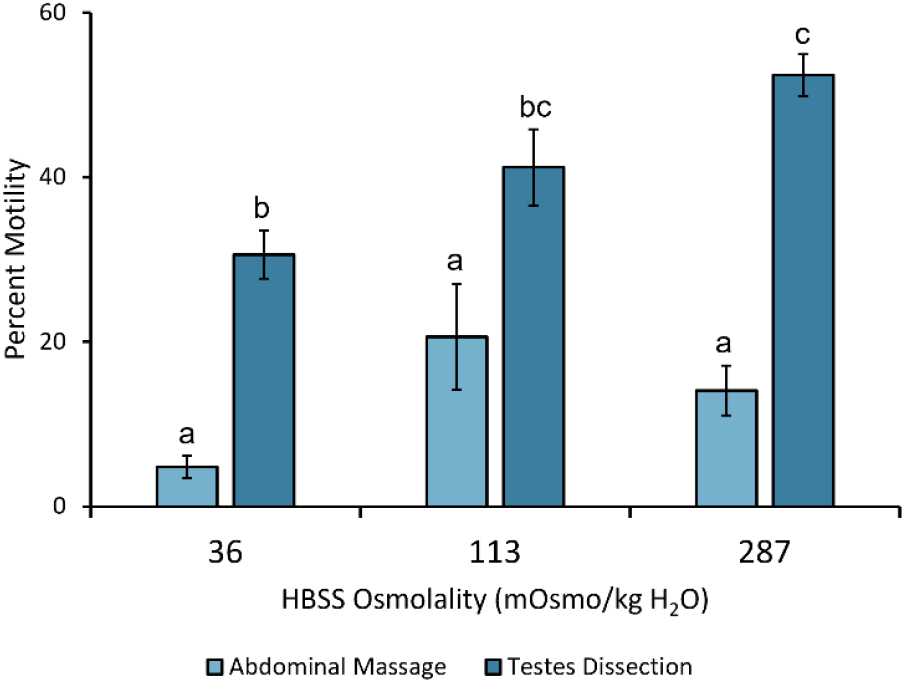
Motility by osmolality of activating solution. Sperm motility activated with HBSS at 36, 113, and 287 mOsmol/kg H_2_O. For 36 and 113, n=4 with 2 fish pooled per sample. For 287, n=9 with 2 fish pooled per sample. Statistical analyses were performed using an ANOVA with Tukey post-hoc test and significant differences are indicated by different letters. The data are represented as mean ± SEM.

#### Sampling methods: abdominal massage vs testes dissection

Sperm sampled by testes dissection are significantly more motile compared to abdominal massage samples from the same fish (**Figure 8A**). Of the motile sperm, a higher percentage are also progressive, and medium or rapid-moving. In sperm sampled by abdominal massage, more motile sperm are slow moving and non-progressive (**Figure 8B-C**).

**Figure 8:**
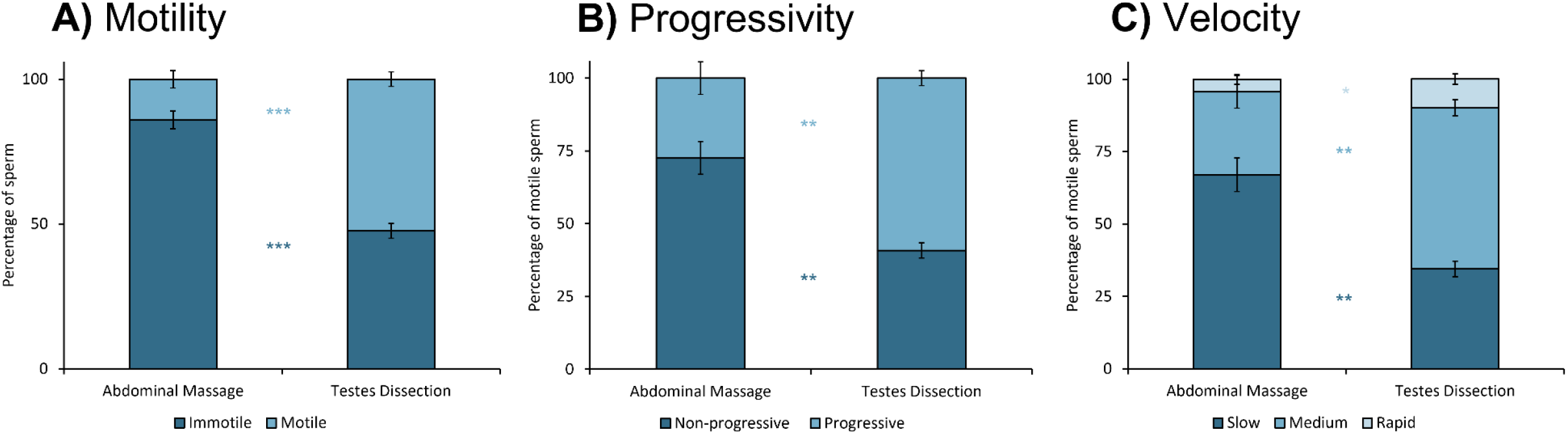
Abdominal massage vs testes dissection. Percentage of **A)** immotile and motile sperm sampled by abdominal massage or testes dissection. Percentage of motile sperm that are **B)** non-progressive and progressive sperm, and **C)** slow, medium, and rapid-moving. All sperm samples were activated in HBSS (287 mOsmol/kg H_2_O). Statistical analyses were performed using a Mann-Whitney U test and significant differences are indicated by asterisks. The data are represented as mean ± SEM, n=9.

#### Motility duration according to storage condition

Sperm sampled by testes dissection and maintained at 27 °C had a 50% reduction in motility in the first 30 minutes after activation. After 2.5 hours, less than 5% of sperm were motile. When stored at 4 °C, motility was reduced by only 14% in the first 30 minutes, and it took 5 hours to see a 50% reduction from the initial motility. Ice had a similar extending effect, but was less effective, with a 26% motility reduction in the first 30 minutes. (**Figure 9A**). Progressivity also dropped 52% in the first 30 minutes on ice, compared to 65% at 27 °C and 33% at 4 °C (**Figure 9B**). Both on ice and at 4 °C, some sperm (<3%) were still moving 42 hours after activation.

**Figure 9:**
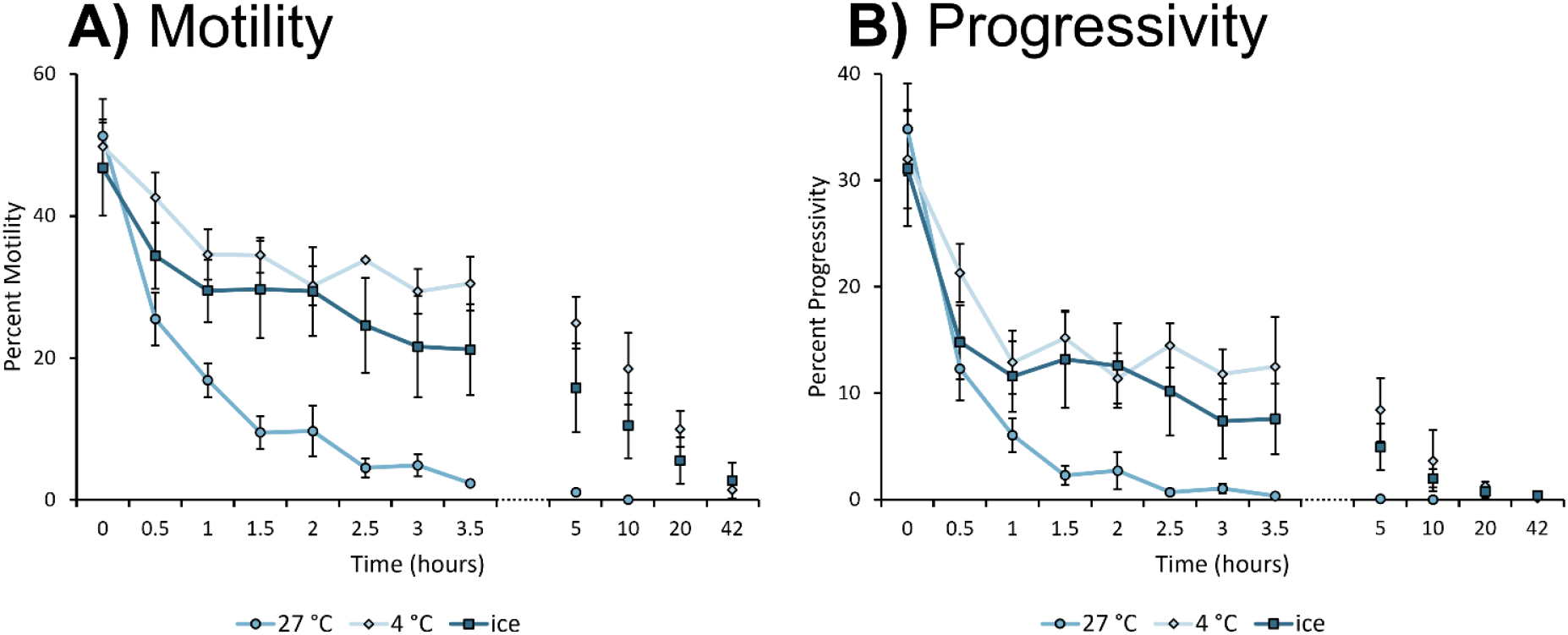
Sperm longevity according to storage condition. Percent **A)** motility and **B)** progressivity over time for samples activated with HBSS (287 mOsmol/kg H_2_O) and stored at 27 degrees, 4 degrees, or on ice. The data points are mean ± SEM, n= ≥4.

#### Importance of the environmental and housing conditions

Males co-housed with females were sampled by testes dissection in the morning before having an opportunity to spawn (before lights turned on), or after the lights turned on and females in the tank had eggs. There was no significant difference in sperm motility. Males housed without females for a month were also sampled, and while there was a trend of higher motility, the difference was also not significant (**Figure 10**).

**Figure 10.**
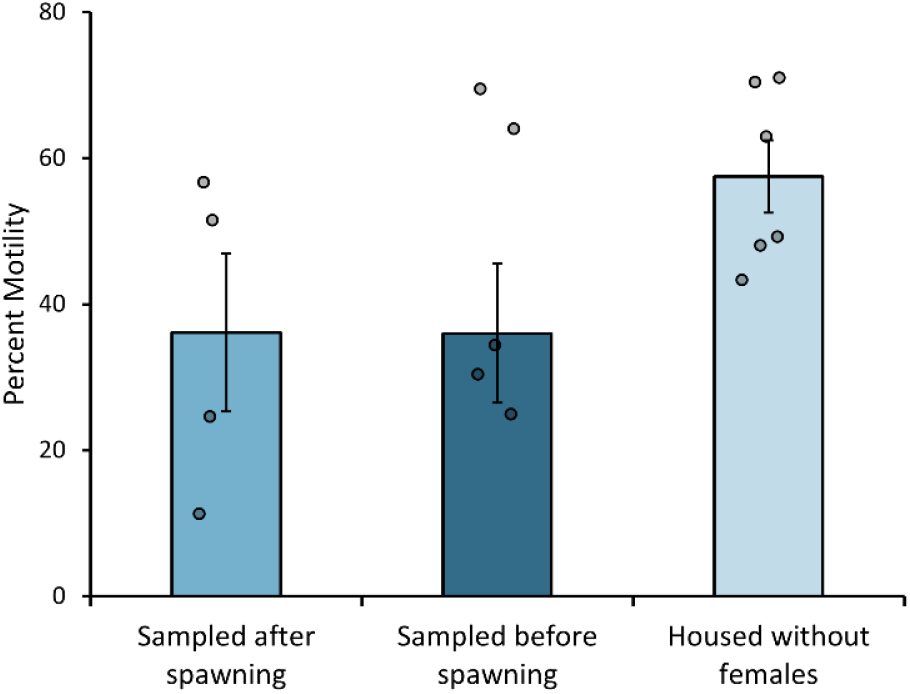
Effect of environmental conditions on sperm motility. Percent motility of sperm sampled before (n=5) and after (n=4) spawning from males that were co-housed with females, and sperm from males housed without females (n=6). All samples were sampled by testes dissection and activated with HBSS (287 mOsmol/kg H_2_O). Statistical analyses were performed using t-tests and differences were not significant. The data is shown as mean ± SEM with circles representing individual fish.

## DISCUSSION

### Activating Solution

Osmolality is an important factor in the activation of fish sperm (Alavi & Cosson, 2006; Wilson-Leedy et al., 2009). Generally, sperm are immotile in the testes and become motile in media that is hyperosmotic relative to seminal fluid for marine fishes, and hypo-osmotic relative to seminal fluid for freshwater fishes (Alavi & Cosson, 2006). Like blood, seminal plasma in freshwater fishes is typically lower than that of marine fishes (about 300 mOsmol/kg compared to 400 mOsmol/kg) (Alavi & Cosson, 2006; Browne et al., 2015). Thus, fish sperm is usually activated on contact with the water in which they live, and this water serves as the best and most biological activation medium for sperm analysis. However, medaka sperm sampled by abdominal massage and testes dissection were not motile in aquarium water (**Figure 6A**), and motility was higher in 287 mOsmol/kg HBSS than in 36 mOsmol/kg HBSS (**Figure 7**), which is unusual for a freshwater fish.

Most previous studies involving sperm analysis in medaka used HBSS (Kowalska A et al., 2007; Kowalska et al., 2011, 2020) or Yamamoto solution (Hara Y et al., 2007; Hashimoto S et al., 2009) as activating mediums for medaka sperm without discussing osmolality. While HBSS is a good activating medium for medaka sperm, variation in osmolality can affect the motility. It is therefore essential for sperm analysis studies to disclose the osmolality of their activating solution for comparison between experiments. One study investigating the ideal osmolality of activating medium found that medaka sperm is motile in deionized water (25 mOsmol/kg) and HBSS with osmolality values <686 mOsmol/kg, with the highest motility between 25 and 227 mOsmol/kg (Yang & Tiersch, 2009). However, in our lab, sperm were not motile in deionized water either. As a euryhaline fish, medaka are very adaptable to different environments and can even live and reproduce in salt water (Inoue & Takei, 2003), so it’s possible that different rearing conditions are responsible for this discrepancy. Interestingly, no other studies have reported testing medaka sperm motility in aquarium water (or similar, like aged tap water), although it is the standard sperm activation medium for freshwater fish.

One possible explanation for this atypical sperm activation in medaka is interaction with ovarian fluid. While fish sperm is usually activated in the surrounding water, ovarian fluid increases or extends the duration of sperm motility in many species (Zadmajid et al., 2019). Although medaka are externally fertilizing freshwater fish, their spawning behavior, which includes fin wrapping and quivering, puts them in much closer proximity than broadcast spawners (Arias Padilla et al., 2021). As the osmolality of ovarian fluid in freshwater fish (about 300 mOsmol/kg) is close to HBSS (Zadmajid et al., 2019), it is possible that fluid released by the female activates the sperm, not the aquarium water. Since the osmolality of seminal and ovarian plasma are similar, it is possible that the ionic composition of medaka ovarian fluid plays an important role (Cosson et al., 2008; Zadmajid et al., 2019). The role of ovarian fluid and ions in activating medaka sperm warrants further investigation.

### Abdominal Massage vs Testes Dissection

In previous studies, medaka sperm has been collected primarily by crushing dissected testes (Kowalska A et al., 2007; Kowalska et al., 2011; Yang & Tiersch, 2009), or by abdominal massage to express the semen directly into activating medium (Hara Y et al., 2007; Hashimoto S et al., 2009; Kawana et al., 2003). Collecting semen by abdominal massage into a capillary tube, a common practice in zebrafish and other teleost fish (Beirão et al., 2009, 2019; Draper BW & Moens CB, 2009), is a viable method in medaka as well. Semen is easily visualized in the capillary tube, and although the volume is too small to be accurately measured, this method enables confirmation of successful collection and analysis of color as a fast indicator of quality (Wasden et al., 2017). Avoidance of fecal contamination is also easy with collection in a capillary tube compared to in medium. Although sperm samples collected by testes dissection have better motility than samples collected by abdominal massage (**Figure 8**), abdominal massage is a minimally invasive procedure that does not require euthanasia and can be repeated in the same fish. Therefore, it is useful for experiments that follow the same fish over time. Also, samples collected with abdominal massage contain only mature cells released with seminal plasma (Beirão et al., 2019), while samples from testes dissection may include immature sperm cells and other debris. The CASA system discounts round cells and debris larger than the max area, but if this parameter is set too high, the motility results could be affected.

Due to the low amount of sperm collected with the abdominal massage technique it was impossible to measure sperm volume and thus to calculate sperm concentrations using regular approaches. However, for samples taken by testes dissection, by keeping the volume of activation medium consistent and weighing the testes, the relative concentration can be calculated based on sperm concentration given by the CASA system to compare between individuals or treatment groups.

### Importance of the sampling conditions

For some species, like zebrafish, it is best to sample sperm before they have a chance to spawn in the morning (Ransom & Zon, 1998) or to isolate males in individual tanks the night before to prevent spawning (Wasden et al., 2017). Our results show that, with medaka, environmental conditions like timing of sampling (before or after spawning) and housing conditions (with or without females) did not have a significant effect on sperm motility (**Figure 10**). It is possible that, with a larger sample size, isolating males from females for a short period of time could yield higher sperm motility, but for experiments where is important to cohabitate both sexes, it does not seem necessary to separate them to yield good results. It also does not seem necessary to sample males at a time before spawning.

### Method of sperm quality analysis

Methods of sperm analysis vary widely for medaka and are often subjective, making results difficult to compare between studies. A study comparing subjective and objective methods used by technicians with varying levels of expertise found that highly experienced technicians can estimate fish sperm motility within 10 percentage points of the data provided by a CASA motility program, while medium and low experience technicians overestimate the CASA motility values with amplitudes up to 30 percentage points (Gallego et al., 2018). However, a lack of standardization for the parameters that determine motility in medaka can also cause variation between those using more objective methods. For instance, one study that recorded sperm at 33 frames per second (fps), analyzed 30 frames, and considered sperm moving faster than 2 µm/s motile had an average velocity of about 60 µm/s and motility of about 70% for their control fish (Hashimoto S et al., 2009). Another study using the same protocol had an average velocity of 40 µm/s for control sperm and a percent motility over 80% (Hara Y et al., 2007). Another group, that analyzed 200 frames at 47 fps, had an average VCL over 100 um/s for control fish, but an average motility below 50% (Kowalska A et al., 2007; Kowalska et al., 2011). They did not disclose what parameters determined motility. So, in this protocol, we use a computer-assisted sperm analysis software to analyze sperm objectively, quickly, and reliably based on a set of parameters that have been customized for the characteristics of medaka sperm. With the complete configuration we used available (**Figure 4B**), this protocol can be reliably replicated in a different lab by different researchers.

### Determining motility parameters

Teleost fish show a wide diversity of sperm characteristics, so although we began with the recommended zebrafish parameters for the SCA Evolution software, it was apparent that the parameters would need to be adjusted for the lower-velocity, greater-longevity medaka sperm. So, we adapted the zebrafish parameters for medaka using literature from medaka and other species that reported similar sperm characteristics (Acosta et al., 2016; Castellini et al., 2011; Hara Y et al., 2007) and selected the thresholds that best fit the distribution of 17,580 analyzed sperm tracks from 18 individuals. Motility is based on curvilinear velocity (VCL) with immotile < 10 µm/s ≤ slow < 20 µm/s ≤ medium ≤ 40 µm/s < rapid. Sperm are considered progressive if straightness index (STR) > 68%. We kept the threshold defining motility at 10 µm/s rather than 2 µm/s, as used in some literature (Hara Y et al., 2007; Hashimoto S et al., 2009), as many sperm lacking flagellar movement were mislabeled as motile with this setting. Other fish species with similar sized sperm heads (∼ 2 µm) also used 10 µm/s to define motile sperm (Acosta et al., 2016; Beirão et al., 2009). We decreased the maximum area from the zebrafish setting of 90 µm^2^ to 20 µm^2^ so large cellular debris from testes dissection would be ignored.

### Motility duration

Duration of motility appears to be dependent on the amount of ATP stored prior to activation, as flagellar movement requires a rapid consumption of energy. Likely due to this rapid exhaustion, there is a correlation between high initial velocity sperm and a shorter duration of motility (Cosson, 2010). While an osmotic shock is typically required to activate fish sperm, it can also lead to membrane damage during the motility period, which can also impact the longevity. This effect is more critical in freshwater species (Cosson, 2010), which may explain why marine sperm are capable of longer motility duration (about 550 sec on average compared to about 150 sec for freshwater species), despite exhibiting a similar average velocity and motility as freshwater sperm (Browne et al., 2015; van der Horst et al., 2021). Motility duration longer than 30 minutes is not common, but it has been reported in several marine species (Browne et al., 2015; Cosson, 2010). Despite being a freshwater fish, our results with medaka fit the profile of sperm that have a lower velocity and longer duration from marine species. This may be related to the activation in medium similar to seminal plasma--without an osmotic shock, medaka sperm likely does not suffer membrane damage like other freshwater fish.

A non-activating solution and very quick, consistently timed analysis after activation is often necessary for analysis of sperm that are only motile for minutes. However, our results show that medaka sperm motility naturally lasts several hours and higher motility may be preserved by storing at 4 °C or on ice (**Figure 9**). While motility does still decrease, because it is gradual, the results are less impacted by minor differences in time of analysis following activation.

## Conclusion

As sperm analysis results are very dependent on the methods used, detailed and reliable methods that can be easily repeated in different labs are beneficial. It is also vital for papers to disclose specifics about their methods to enable repeatability. With medaka used increasingly as a model for reproduction research, the body of information regarding assessment of sperm quality is lacking. The two distinct methods described in this paper for sperm sampling may be beneficial for different experiments. Testes dissection yields higher motility sperm and allows for relative concentration calculations, while abdominal massage can be done repeatedly on the same fish and is a more pure, biological representation of a spawning event. We therefore provide the parameters for CASA, which is reliable, objective technique that provides ample data about sperm movement, including motility, progressivity, velocity, and other kinematic parameters. These protocols will thus be useful for a variety of studies in medaka including toxicology, ecology, reproduction, and physiology.

## ACKNOWLEDGMENTS

This work has been funded by the Norwegian University of Life Science and the U.S. Fulbright program.

